# Epistasis-mediated compensatory evolution in a fitness landscape with adaptational tradeoffs

**DOI:** 10.1101/2024.11.03.621663

**Authors:** Suman G. Das, Muhittin Mungan, Joachim Krug

## Abstract

The evolutionary adaptation of an organism to a stressful environment often comes at the cost of reduced fitness. For example, resistance to antimicrobial drugs frequently reduces growth rate in the drug-free environment. This cost can be compensated without loss in resistance by mutations at secondary sites when the organism evolves again in the stress-free environment. Here we analytically and numerically study evolution on a simple model fitness landscape to show that compensatory evolution can occur even in the presence of the stress and without the need for mutations at secondary sites. Fitness in the model depends on two phenotypes – the null-fitness defined as the fitness in the absence of stress, and the resistance level to the stress. Mutations universally exhibit antagonistic pleiotropy between the two phenotypes, that is they increase resistance while decreasing the null-fitness. Initial adaptation in this model occurs in a smooth region of the landscape with a rapid accumulation of stress resistance mutations and a concurrent decrease in the null-fitness. This is followed by a second, slower phase exhibiting partial recovery of the null-fitness. The second phase occurs on the rugged part of the landscape and involves the exchange of high-cost resistance mutations for low-cost ones. This process, which we call *exchange compensation*, is the result of changing epistatic interactions in the genotype as evolution progresses. The model provides general lessons about the tempo and mode of evolution under universal antagonistic pleiotropy with specific implications for drug resistance evolution.

---

Compensatory evolution occurs when the deleterious effects of mutations on a phenotype or fitness are partly or fully reversed through subsequent evolution [1–7]. The initial deterioration can occur when deleterious mutations are fixed through drift or hitchhike on beneficial mutations [2]. Alternatively, the adaptation of an organism to a challenging environment can lead to selection for beneficial mutations that are detrimental in the original environment. Evolutionary mechanisms can eventually compensate for this loss without affecting the fitness gain in the new environment. This is the case we study in this paper.

A classic example of this kind of compensatory evolution is provided by drug resistance, where resistance mutations selected upon exposure to a drug often reduce the *null-fitness* (defined as fitness in the original environment, *i*.*e* in the absence of the drug; see [8–11] and references therein). A potential consequence of this is that bacteria selected for high resistance grow more slowly, which affects their transmission to new hosts and reduces their ability to cause widespread infection. However, experimental studies in which resistant microbes are cultured in the absence of drugs find that these evolve to (partially) regain their null-fitness [4, 9, 12, 13]. The most commonly reported mode for this process is through compensatory mutations at secondary sites that do not compromise the acquired resistance [4, 9, 14, 15], though the contribution of the reversion of resistance mutations has been noted in some cases as well [3, 4, 12, 16]. Reduction of the cost of resistance can also happen in the presence of the drug [12, 17–20] However, only a small number of empirical studies have directly addressed this question [14, 21, 22], despite its importance in determining the optimal course of drug treatment [12]. It has been found that the compensation loci and fitness effects in the presence of drugs can be different from those in its absence [14, 21], but due to the limited literature on the subject, there is no consensus on general mechanisms of compensatory evolution in the presence of drugs, or indeed on how generic it is.

In this theoretical work, we present modeling results that demonstrate a mode of compensatory evolution that does not require reversion to the original environment and mutations at secondary sites. To do this, we focus on how the structure of the fitness landscape (which captures the effect of gene-gene interactions, that is epistasis, on fitness) guides the evolution of resistance to an environmental stressor. A number of studies have constructed combinatorially complete landscapes comprising a small number of loci [23–28] but our understanding of resistance evolution on large fitness landscapes and how it is impacted by the environment remains incomplete. Here we study evolution on an empirically-motivated fitness landscape model where every resistance-increasing mutation also reduces the null-fitness. Epistasis is introduced by the coupling of the two phenotypes, namely null-fitness and resistance, in producing the net fitness.

We find that when the wild type is subjected to a fixed stress level, evolution is biphasic. The first phase exhibits a gain in resistance accompanied by a loss in null-fitness, and the second phase shows a substantial amount of compensatory evolution of null-fitness even as the stress parameter stays constant. Compensation occurs through the reversal of high-cost resistance mutations and the substitution of low-cost ones. Contrary to common wisdom that regards compensatory mutations as a separate class of events, we show that they are an emergent feature of a model that contains only one class of mutations (namely, mutations with antagonistic pleiotropic effects [29] on resistance and null-fitness). It is the changing selection pressure along an evolutionary path that singles out mutations with qualitatively different phenotypic effects in the two phases of adaptation.

## I. MODEL AND TERMINOLOGY

### A. Fitness landscape model

We focus on an empirically-grounded model of tradeoff-induced landscapes (TIL model, see [30, 31]) to study the evolution of of a haploid population exposed to environmental stress. While we use examples from the well-developed literature on drug resistance for illustration purposes, our results are primarily about a new and generic mechanism for compensatory evolution. For a given genotype ***σ***, the two relevant phenotypes are the stress resistance level *m*_***σ***_ and the null-fitness denoted by *r*_***σ***_. Resistance comes at a cost, in the sense that resistance-increasing mutations reduce the null-fitness. The fitness of a genotype ***σ*** as a function of an environmental stress variable (such as drug concentration) *x* is represented by the population growth rate, which is assumed to be given by a response curve of Hill type [32],

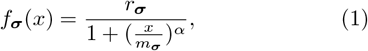

where *α >* 0 is the Hill coefficient. The parameter *x* represents an environmental challenge that reduces the fitness of all genotypes. Response curves of different genotypes intersect as *x* changes, thus altering the rank order in the landscape; see supplementary Fig. S1. (For further graphic illustrations of dose-response curves or DRCs for antimicrobial drugs and their relevance to fitness landscapes, see [30, 31, 33].)

The genotype is encoded by a binary vector ***σ*** = (*σ*_1_, …, *σ*_*L*_), where *L* is the number of mutation sites of interest and *σ*_*i*_ = 0, 1 indicates the absence or presence of a mutation at locus *i*. We assume that the mutational effects on null-fitness and resistance are non-epistatic (multiplicative), that is

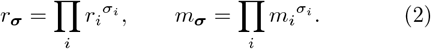

As we will see in Section 3C, this choice makes the model more analytically tractable, yet retains the features of adaptive evolution that we are interested in. Moreover, some experimental studies of antibiotic resistance evolution in bacteria show that Eq. (2) holds approximately [30, 34]. (See [30] for more details on the empirical motivation behind the model and its relevance.) It is convenient to work with log-transformed parameters *u*_*i*_ =− ln *r*_*i*_, *v*_*i*_ = ln *m*_*i*_, *𝒳* = ln *x, F*_***σ***_ = ln *f*_*σ*_. With this definition,

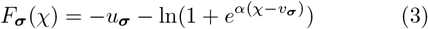

where *u*_***σ***_ = ∑ *σ*_*i*_*u*_*i*_, *v*_***σ***_ =∑ *σ*_*i*_*v*_*i*_ define the additive genotype-phenotype maps of *u* and *v*. The assumed tradeoff between resistance and null-fitness implies that *r*_*i*_ *<* 1, *m*_*i*_ *>* 1 and therefore *u*_*i*_ *>* 0, *v*_*i*_ *>* 0 for all *i*. Further, we impose the condition that

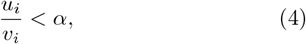

because, as shown later, any mutation violating this condition is always deleterious, and can thus be ignored. The TIL model assumes universal antagonistic pleiotropy [29], in that every mutation affects both null-fitness and resistance in opposite directions. To generate large fitness landscapes, the {*u*_*i*_} and {*v*_*i*_} are taken to be random variables, where each pair (*u*_*i*_, *v*_*i*_) is drawn independently from a distribution *P* (*u*_*i*_, *v*_*i*_) which respects the constraints imposed by the tradeoff and by Eq. (4). We denote 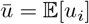 and 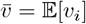, where 𝔼 [·] is the mean with respect to the joint distribution of *u*_*i*_ and *v*_*i*_. Note that the TIL model has variable tradeoffs, that is, the value of the resistance benefit *v*_*i*_ does not uniquely determine the cost *u*_*i*_.

### B. Evolutionary dynamics

We are interested chiefly in evolution at fixed values of the parameter *𝒳*. We focus initially on the strong selection/weak mutation (SSWM) regime where selection is sufficiently strong so that only beneficial mutations can fix in the population, and mutations are sufficiently rare so that a new mutation occurs only after the previous mutation has been fixed or removed from the population [35]. Later, in Section 2D we will relax this assumption. In the SSWM limit, the evolutionary dynamics is an adaptive walk, in which the population moves along the fitness landscape in single mutational steps that each increase fitness. The walk terminates when the population reaches a local fitness peak, *i*.*e*. a genotype which has higher fitness than all other genotypes that are one mutation away. The probability that a *de novo* mutant ***σ***′ occurring in a population with genotype ***σ*** is fixed is given by the large population-size limit of the Kimura formula, 1 − *e*^−2*s*^ with the selection coefficient *s* max 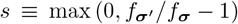. We denote an adaptive walk with this fixation probability as the *Kimura adaptive walk*. We will also consider the *uniform adaptive walk*, where every fitter mutant has equal probability of being fixed [36]. A single time step of the adaptive walk consists of the origin and fate (fixation or extinction) of a mutation [37]. Therefore, the number of time steps is equal to the total number of mutations that have occurred (but not necessarily fixed).

## II. RESULTS

We will first describe the broad features of the TIL fitness landscapes and the adaptive walks based on simulations. We then turn to a qualitative explanation of the adaptive evolution at a phenotypic level. Finally, we show that the results described in the first two subsections can be derived in precise mathematical terms, if we neglect the fluctuations in resistance by assuming that each mutation contributes a fixed amount *v* to the log-resistance *v*_***σ***_ of a genotype.

### A. Directed and fluctuating phases of adaptation

The structure of the fitness landscape and an adaptive walk trajectory starting at the wild type are illustrated schematically in Fig. 1. Since both the null-fitness and the resistance contribute to the microbial fitness but there is a tradeoff between the two, fitness maxima are expected to have an intermediate number of mutations, the distribution of which is governed by *χ*. For large *L* and for *χ* ~ *O*(*L*), the leading order approximation to the mean number of mutations *q*\* in a fitness peak is given by 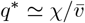 (see [30] and SI text). The fitness peaks are located in a narrow band situated around *q*\* (Fig. 1).

**Fig 1.**
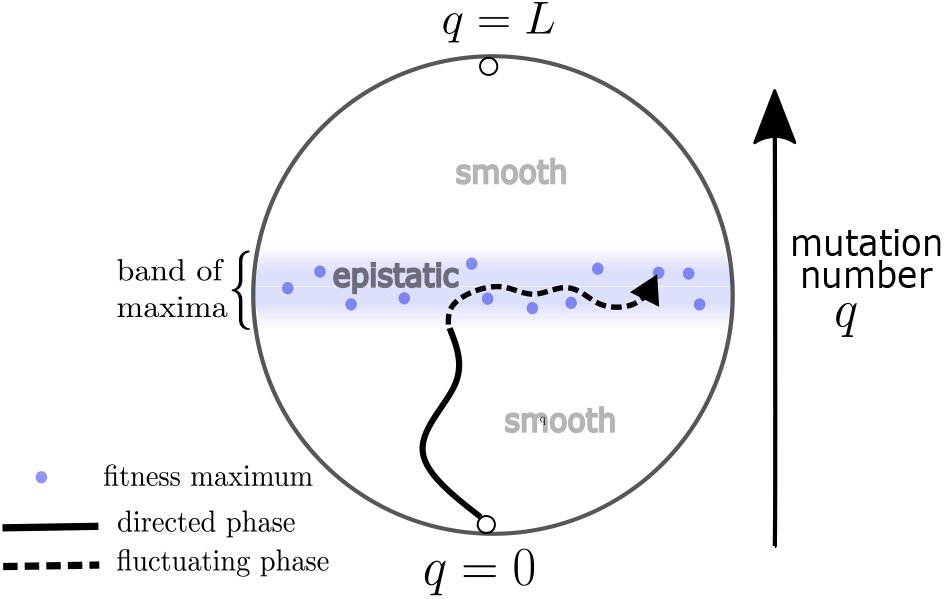
The figure depicts a schematic for visualizing the TIL model and adaptive walks on it. The genotypes are depicted as inhabiting the circle, and the number of mutations in the genotypes increases in the vertical direction. At any given stress parameter *𝒳*, the local fitness maxima (depicted as filled blue circles) contain a typical number of mutations, and therefore lie in a narrow band shaded in blue. An adaptive walk trajectory is shown as a black line. The walk begins at the wild type (*q* = 0) and terminates at one of the maxima. The trajectory has two phases. The solid part of the black line shows the directed phase, where the mutation number grows monotonically. The trajectory transitions to the fluctuating phase, marked with the dashed part of the black line, once the band of maxima is reached.

An adaptive walk starting from the wild type has two distinct phases. The first is the *directed* phase, where the mutation number *q* in the evolving genotype increases monotonically (Fig. 2A). The term ‘directed’ refers to the fact that all fitness-increasing mutations in this phase are of the form *σ*_*i*_ = 0 → 1, *i*.*e*. there is no reversal of acquired mutations and mutations accumulate. Naturally, this phase is associated with a monotonic increase in resistance along with a monotonic decrease in null-fitness, as shown in Fig. 2B and C. As depicted in Fig. 1, the directed phase ends once the walk reaches the narrow band of maxima. The system now enters the *fluctuating* phase, where the mutation number can go down as well as up, and the dynamics includes both forward (0 → 1) and reverse (1 → 0) mutations. In the fluctuating phase, the mean resistance is nearly constant whereas the mean null-fitness increases, leading to a partial recovery of the cost of resistance. Figure 2C shows that the recovery is a slow process. This is partly due to the fact that there are fewer beneficial mutations, but also because there is a drop in the selection coefficients and therefore in the fixation probability of beneficial mutations, as seen in Fig. 2D. In fact, it is shown in SI Text and Fig. S2 that for large *L* the fixation probability undergoes a *sharp* transition between the two phases, in the sense that the time-scale over which the transition occurs is much smaller than the time-scale for the walk.

**Fig 2.**
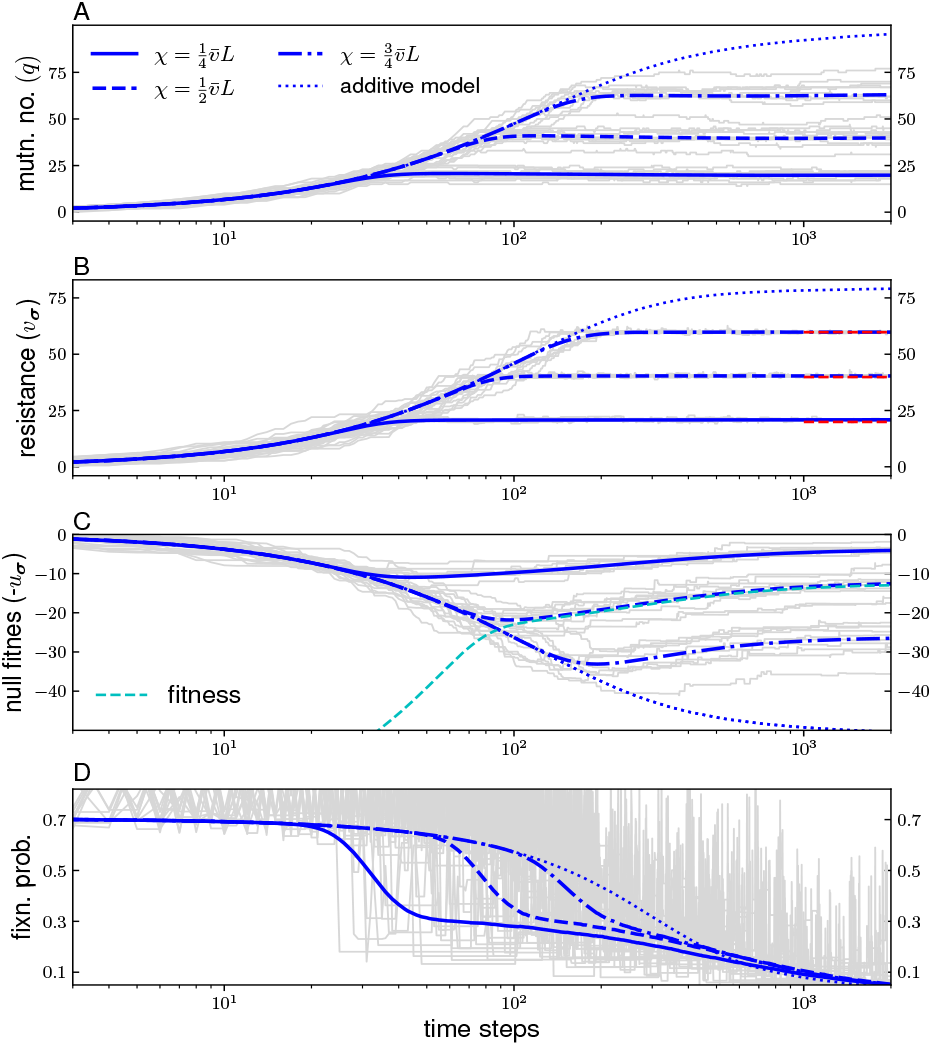
Numerical results for the mean of the various quantities associated with the evolving genotype are plotted. The mutational effects (*u*_*i*_, *v*_*i*_) are chosen from the distribution given in the Methods section, with *γ* = 0. The genome size is *L* = 100. The highest relevant value of *𝒳* is 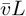, which is the mean *v*_***σ***_ of the most resistant mutant. We have chosen three equally spaced *𝒳* values below this threshold to show how evolution depends on the stress level. The colored lines are computed by generating a random landscape, simulating a Kimura adaptive walk, and averaging over 10^4^ such realizations, while gray lines show individual trajectories. The dotted lines are averages for the additive model described by Eq. (5). (A) The mean mutation number and individual trajectories are shown for 3 different values of *𝒳* as indicated in the panel. The same values are used in the remaining panels. (B) The mean resistance level saturates to a value close to *𝒳* (dashed red lines) in the fluctuating phase. (C) The null-fitness decreases in the directed phase and then grows in the fluctuating phase through the process of exchange compensation (see main text). The cyan line shows the fitness evolution for 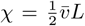. The fitness closely follows the null-fitness as the resistance saturates to *v*_***σ***_ ≃ *𝒳* in the fluctuating phase. (D) The fixation probability of all beneficial single-mutant genotypic neighbors of the evolving genotype are shown.

The origin of the directed phase can be understood as follows. Equation 3 together with Eq. (4) implies that when 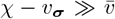 (or equivalently *q*\* − *q*_***σ***_ ≫1), every forward mutation increases the fitness. In this phase, the fitness given by Eq. (3) can be approximated as *F* ≃− *u*_***σ***_ + *αv*_***σ***_− *α𝒳*. The constant term *αχ* has no effect on the dynamics and can be dropped, leading to an additive version of the TIL model,

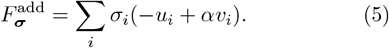

Thus, the system starting at the wild type (*q*_***σ***_ = 0 and *v*_***σ***_ = 0) traverses an effectively smooth landscape given by Eq. (5), acquiring mutations sequentially until its resistance reaches the value *v*_***σ***_ ≃ *𝒳* (Fig. 2B). The departure of the full TIL model from the additive model indicates the transition from the directed to the fluctuating phase, as seen in Fig 2.

The fluctuating phase exhibits compensatory evolution of the null-fitness. There are two notable features of this phase. Firstly, it demonstrates the role of epistasis in shaping evolutionary trajectories in this model. The fluctuating phase is a combination of forward and backward mutations, where the latter exhibits reversal of some of the resistance mutations acquired during the directed phase. Clearly, in the course of adaptive evolution, the effect of these mutations has switched from being beneficial (in the directed phase) to deleterious (in the fluctuating phase), exhibiting sign epistasis [5]. Although the genotype-phenotype map is additive, epistasis in fitness is introduced by the non-linear dependence of fitness *F* on the resistance *v* in Eq. (3), as noted in [30]. In the terminology of [38] this constitutes an example of non-specific epistasis. Secondly, we point out that compensatory evolution in this model occurs in the presence of universal antagonistic pleiotropy, in contrast to standard modes of compensatory evolution reported in the literature. Indeed, every reversal of a resistance mutation not only increases the null-fitness but also reduces the resistance. The role of pleiotropy in guiding the evolution of the phenotypes *u* and *v* is discussed in more detail in the next section.

**TABLE 1.** List of main mathematical terms.

|  |  |
| --- | --- |
| $L$ | number of loci in a genotype |
| $\sigma$ | genotype (expressed as binary sequence of length $L$) |
| $\sigma^{+i}$ | genotype obtained from $\sigma$ by addition of a mutation at site $i$ |
| $\sigma^{-j}$ | genotype obtained from $\sigma$ by removal of a mutation at site $j$ |
| $\alpha$ | Hill coefficient (indicates the steepness of fitness response to environmental stress) |
| $u_\sigma$ | fitness cost of genotype $\sigma$ |
| $v_\sigma$ | stress resistance level of genotype $\sigma$ |
| $F_\sigma$ | fitness of genotype $\sigma$ |
| $q$ | number of mutated loci in the genotype |
| $q^*$ | number of mutated loci in a fitness peak |
| $\chi$ | value of environmental stress variable |
| $\chi_i$ | value of stress variable beyond which the single mutant with mutation at locus $i$ is fitter than the WT |
| $\chi_{\sigma,+i}$ | value of $\chi$ beyond which the genotype $\sigma$ with additional mutation at locus $i$ is fitter than $\sigma$ |
| $c_i$ | relative cost of a resistance mutation |
| $\gamma$ | parameter that controls tradeoff between null-fitness and resistance |
| $I^+$ | set of all mutated loci in a genotype |
| $K(\chi)$ | the number of single mutants which at $\chi$ are fitter than the WT |
| $\kappa$ | difference between largest and smallest values of $\chi$ at which single mutants become fitter than WT |
| $\mu$ | mutation rate per individual per generation in Wright-Fisher simulations |
| $N$ | population size in Wright-Fisher simulations |

### B. Trajectories in phenotypic space and the role of pleiotropy

Several features of this model can be qualitatively understood purely at the phenotypic level by observing that (i) the response curve given by Eq. (3) furnishes a phenotype-fitness map, and (ii) evolution in the two-dimensional phenotypic space (*u*_***σ***_, *v*_***σ***_) is constrained by universal antagonistic pleiotropy, represented by the condition *u*_*i*_, *v*_*i*_ *>* 0. A phenotypic mutation (*u*_***σ***_, *v*_***σ***_) → (*u*_***σ***_ +*u*_*i*_, *v*_***σ***_ +*v*_*i*_) induces a fitness change 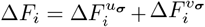, where

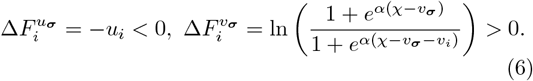

The quantity 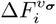 is monotonic increasing in *χ* and reaches the limit *αv*_*i*_ as *𝒳* → ∞. Thus, for a mutation with *u*_*i*_ *> αv*_*i*_, the fitness change Δ*F*_*i*_ is negative for all *χ*. Consequently, such a mutation will never be fixed, as noted before in Eq. (4). This also implies that the reversal of a mutation violating Eq. (4) is always beneficial, and it will eventually be lost if it happens to be initially present in the genotype.

The phenotype-fitness map implies that a mutant that is fitter than the wild type at *χ* must satisfy the condition

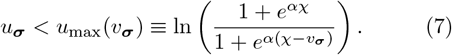

The function *u*_max_(*v*_***σ***_) is the maximum cost that can be incurred in order to acquire resistance *v*_***σ***_, and further, *u*_***σ***_ *< u*_max_(*v*_***σ***_) delimits the region of the phenotypic space where adaptive evolution is possible. This is shown as the green shaded region in Fig 3A.

**Fig 3.**
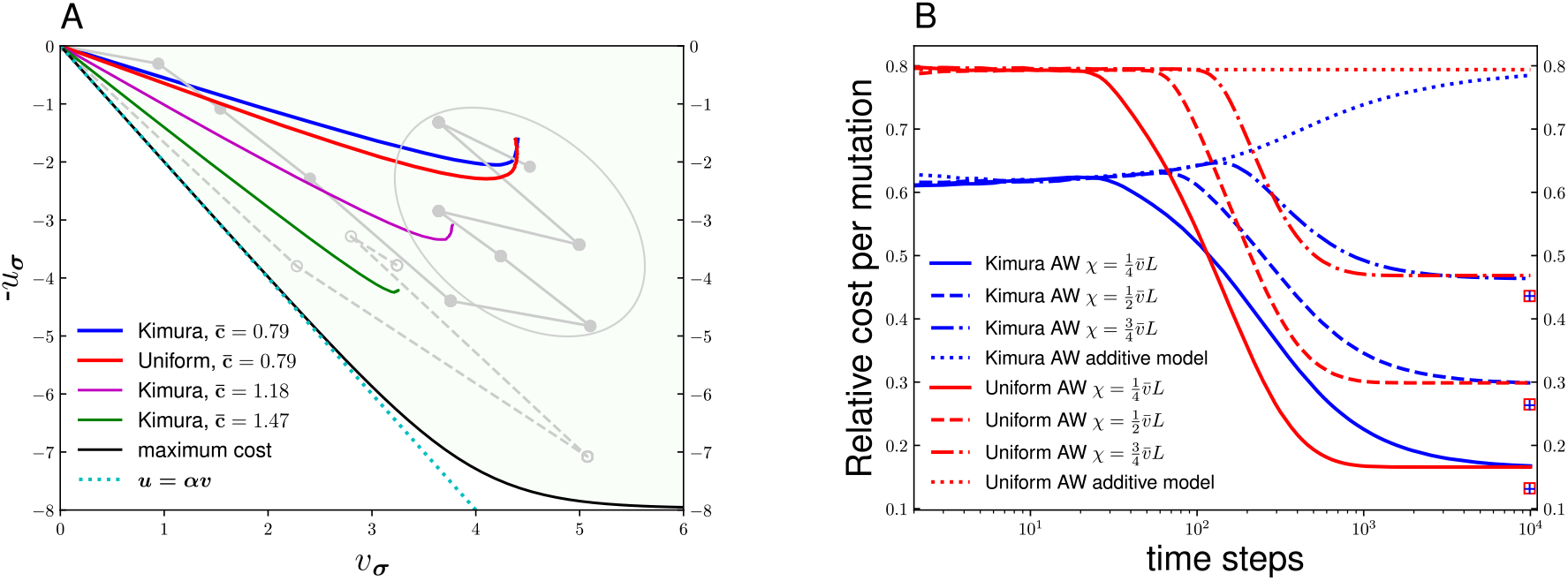
**A**. The plot shows the evolutionary paths in the *u* − *v* plane for a genotype with *L* = 10 loci evolving at 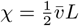. The colored solid lines are averages over 10^4^ realizations. The constant 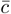 represents the mean relative cost of resistance mutations in the landscape. The distribution of the {*u*_*i*_, *v*_*i*_} is given in Methods. The parameter *γ* is varied to tune 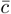 (*γ* = 0 corresponds to 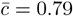). The black line represents the upper bound of the net cost as given in Eq. (6). The green shaded region is the part of the phenotypic space to which adaptive walks are restricted. The curve *u* = *αv* (dotted cyan line) eventually deviates from the line of maximum cost since costly resistance mutations are no longer beneficial once a sufficiently high resistance level is achieved. Also shown are two individual trajectories at low cost (solid grey line) and high cost (dashed grey line). **B**. The plot shows the mean relative cost of the mutations present in the evolving genotype averaged over 10^4^ realizations. The system size is *L* = 100 and all other parameters are the same as in Fig 2. Results are shown both for Kimura and uniform adaptive walks. The dotted lines represent the same quantity in the case of the additive model; the absence of compensation here is because lack of epistasis rules out reversals of mutations. The plus signs and squares represent the theoretical lower bounds averaged over 10^4^ landscape realizations for the Kimura and uniform cases respectively.

The role of pleiotropy is illustrated by the shape of the adaptive walk trajectories in the *u* − *v* plane, plotted in Fig 3A (colored solid lines). In the directed phase, the trajectories become increasingly aligned with the direction of the maximum cost line (black curve) for systems with higher *relative cost* of resistance, characterized by 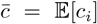, where *c*_*i*_ = *u*_*i*_*/v*_*i*_ is the relative cost of an individual mutation. In the fluctuating phase, there is a gain in mean null-fitness without loss of mean resistance, which may seem puzzling in the presence of antagonistic pleiotropy. The mechanism behind this becomes clearer upon examining specific sample trajectories (gray lines in Fig 3A). By looking closely at the individual adaptive steps (for example, those in the circle enclosing the fluctuating gray line in Fig 3A), we see that the null-fitness increases overall because the gain due to the reversal of resistance mutations is higher than the loss that occurs due to the forward mutations. Meanwhile, the resistance does not change substantially because the forward mutations occurring in the fluctuating phase counter the negative effect of the reversals on resistance. The model allows variable tradeoffs between *u* and *v*, implying that mutations conferring similar levels of resistance may differ considerably in their negative effects on null-fitness. The process proceeds by reversals that occur preferably on resistance mutations that have higher values of *u*_*i*_ and forward mutations that have lower values of *u*_*i*_, without substantial alteration of *v* in the long run. This is treated in greater analytical detail in the next subsection through a special case of the TIL model where all mutations have the same *v*_*i*_. We introduce the term *exchange compensation* to encapsulate the idea that in the fluctuating phase, compensation occurs by ‘exchanging’ acquired resistance mutations bearing a high cost for those with a low cost through a multi-step process. To put it succinctly, *the fluctuating phase selects against the strength of antagonistic pleiotropy while maintaining the resistance level*.

The effect of pleiotropy can be quantified through the evolution of the relative cost *c*_*i*_ associated with the substitutions. As noted before, *c*_*i*_ is bounded above by *α* (Eq. (4)), and bounded below by 0 (corresponding to a non-pleiotropic resistance mutation). Figure 3B summarizes the evolution of the mean relative cost of resistance mutations contained in the genotype, defined as (∑_*i*_ *σ*_*i*_*c*_*i*_) */* (∑ _*i*_ *σ*_*i*_), averaged over several landscape and walk realizations, and conditioned on at least one mutation being present in ***σ***. The results for the additive model in Eq. (5) are plotted as well. The points at which the additive model results diverge from the full TIL model indicate the end of the directed phase. As seen in the figure, the relative cost is high in the directed phase, and goes down as compensatory evolution occurs, reflecting the decreasing pleiotropic strength of the resistance mutations accumulating in the genotype. For the uniform adaptive walk, the mean relative cost in the directed phase (as given by the dotted red line) agrees with the expected relative cost of mutations in the full landscape, 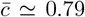, since in the uniform walk all forward mutations are equally likely to be fixed in the directed phase. The Kimura walk behaves in a different way in the directed phase - there is a slight initial decrease since selection prefers both high *v*_*i*_ and low *u*_*i*_, followed by a modest increase as low-cost resistance mutations become rarer. The maximum relative cost accrued by the Kimura walk (which occurs at the end of the directed phase) is lower than that of the uniform walk, which is expected since the former preferably fixes high-fitness mutants. The fluctuating phase of both the Kimura and uniform walks terminate with similar average values of the relative cost.

A lower bound on the per-mutation relative cost of a resistant genotype with *n* mutations is given by the mean of the *n* lowest *c*_*i*_ values from the pool of *L* mutations in the landscape. It is seen in Fig. 3B that the actual relative cost at the end of adaptive walks is not much higher than this bound, indicating that, within the constraints set by the number of resistance mutations, most of the cost is actually recovered through the process of compensation. Note that the degree of recovery for *L* = 100 in Fig. 3B is much higher than for the corresponding *L* = 10 case in Fig. 3A. Indeed, the degree of compensation increases with *L*, as shown in Fig. S3, because for larger *L* the landscape contains mutations with values of *u*_*i*_ that extend closer to zero.

### C. Mechanism of exchange compensation in a simplified setting

In the previous two subsections we have characterized the features of the biphasic adaptive evolution on TIL fitness landscapes. In order to understand better how this evolution emerges from the microscopic dynamics of single fitness increasing mutations, we now turn to a theoretical analysis, characterizing the topography of the landscape.

For a given genotype ***σ*** = (*σ*_1_, …, *σ*_*L*_), we define *I*^+^ = {*i* : *σ*_*i*_ = 1} as the set of loci at which there is a mutation. The complement *I*^−^ of this set comprises the loci where mutations are absent. In what follows we will specify a genotype ***σ*** in terms of its mutation set *I*^+^ [30, 31]. For *i*∈ *I*^−^ we denote the genotype obtained from ***σ*** by adding a mutation at locus *i* as ***σ***^+*i*^, likewise for *j* ∈ *I*^+^, ***σ***^−*j*^ is the genotype obtained from ***σ*** by reverting the mutation at locus *j*.

The response curves associated with a pair of genotypes ***σ*** and ***σ***^+*i*^, differing by a mutation at a single locus, are such that they intersect precisely once, and we denote the value of stress at that point as *χ*_***σ***,+*i*_ (see SI for necessary and sufficient conditions that ensure this). For *𝒳 > 𝒳*_***σ***,+*i*_ the genotype ***σ***^+*i*^ will have larger fitness than ***σ***, so that adding the mutation at *i* is fitness increasing. It is found that

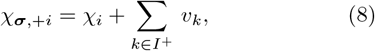

where *χ*_*i*_ is the stress parameter value at which the response curves of the wild-type **0** and the single-mutant **0**^+*i*^ intersect. Thus all intersection points *𝒳*_***σ***,+*i*_ can be constructed from the set of *χ*_*i*_’s and log-resistances *v*_***σ***_. This is a direct consequence of the multiplicative nature of the parametrization in Eq. (2). We label the loci *i* = 1, 2, …, *L* such that

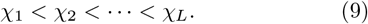

It follows from Eq. (8) that if *χ* is sufficiently large, the addition of any mutation will be fitness increasing. This characterizes the directed phase.

Figure 2B indicates that in the fluctuating phase the statistical fluctuations in the resistance are far less than in the preceding directed phase. This suggests approximating the resistance 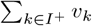 of a genotype by its mean *qv*, where *v* is the average of *v*_*i*_ and *q* = | *I*^+^| is the mutation number. In this approximation, each mutation contributes an equal “quantum” *v* to the resistance of a genotype. We shall call this the *q*-TIL model. This approximation renders the model analytically tractable, while retaining the key features of the adaptive evolution.

The genotype ***σ*** is a fitness peak at *χ*, if for each *i* ∈ *I*^−^ and *j* ∈*I*^+^, the genotypes ***σ***^+*i*^ and ***σ***^−*j*^ have lower fitness. Fitness peaks of the TIL model can be explicitly identified [31]. In the *q*-TIL model, this procedure simplifies and the condition for fitness peaks at *𝒳* becomes^1^

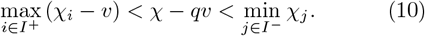

Letting

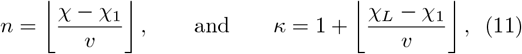

where ⌊*a*⌋ is the greatest integer less than or equal to *a*, and defining

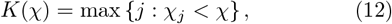

one can deduce from Eq. (10) the following results:

i. in terms of mutation number *q*, the fluctuating region is a strip of width *κ*, given by *n* − *κ* + 1 ≤ *q* ≤ *n* + 1, as shown in Fig. 4A,
ii. all fitness peaks at a given *𝒳* have the same mutation number *q*\*, where *q*\* is the unique choice for *q* such that the following inequality holds:

**Fig 4.**
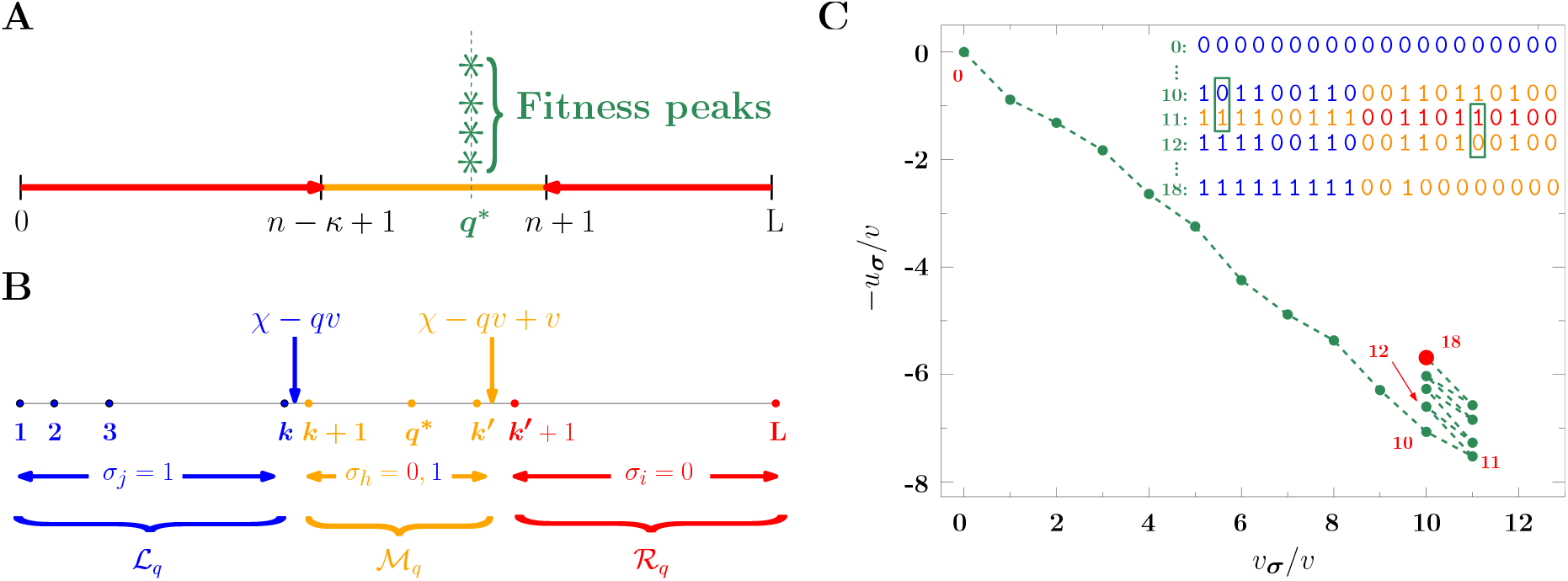
**The** *q***-TIL model, a simplified version of the TIL model where each mutation confers the same resistance** A: All fitness peaks have the same mutation number *q*\* and the directed and fluctuating phases of evolution are determined by the range of mutation numbers indicated in red and orange respectively. In the directed phase (red) the arrows point in the direction of increasing fitness and thus move genotypes into the fluctuating regime (orange) which is trapping. B: The log-stress value *𝒳* together with the mutation number *q* of a genotype partition the set of loci into the three sets marked as *ℒ*_*q*_, *𝓂*_*q*_, and *ℛ*_*q*_. For loci in *ℒ*_*q*_ (*ℛ*_*q*_) adding (removing) a mutation is always fitness increasing. C: Adaptive evolution in the (*u*_***σ***_, *v*_***σ***_) plane. Shown is a realization with *L* = 20 loci in the special case *κ* = 1. Step numbers are shown in red next to selected data points. Up to step 10, the adaptive walk is in the directed phase, where mutations pile up. In the fluctuating phase, the effective dynamics consists of *exchanging* mutation states of pairs of loci. The inset shows genotypic configurations at certain steps of the walk. Refer to text for further details.

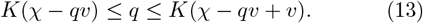

ii. The derivation of these results and further details on the *q*-TIL model will be presented elsewhere. The underlying key observation is that given *𝒳* and *q*, the intersection points of Eq. (8) become *𝒳*_***σ***,+*i*_ = *χ*_*i*_ + *qv* in the *q*-TIL model and in turn partition the set of loci into three subsets *ℒ*_*q*_ = {1, …, *k*}, *𝓂*_*q*_ = {*k* + 1, …, *k*′} and *ℛ*_*q*_ = {*k*′ + 1, …, *L*}, where *k* = *K*(*χ* − *qv*) and *k*′ = *K*(*𝒳* − *qv* + 1). For any locus *i* ∈ *ℒ*_*q*_ (*j* ∈ *ℛ*_*q*_) a transition is fitness increasing if and only if it involves adding (removing) a mutation at that locus, as illustrated in Fig.4B.

In the *q*-TIL model, the bi-phasic adaptation takes a particularly simple form when the width *κ* of the fluctuating region takes its smallest possible value *κ* = 1, so that the genotypes oscillate between *q* = *n* and *q* = *n* + 1. This is illustrated in Fig.4C, showing the evolution of *u*_***σ***_ and *v*_***σ***_ for *L* = 20. The choice of *χ* was such that *n* = *q*\* = 10 and *k* = *K*(*𝒳* − *q*\**v*) = 9.

Starting with the wild-type, the directed phase is characterized by mutation numbers *q* ≤ *n*, cf. Fig.4A. In this regime, the set *ℒ*_*q*_ is the set of all loci and therefore any mutation 0 → 1 increases fitness. As long as *q < n*, mutations are added one by one and as a result the log null-fitness −*u*_***σ***_ decreases monotonically, while the resistance *v*_***σ***_ = *qv* increases. Next, the walk enters the fluctuating phase at step 10, i.e. when *q* = *n* = 10, cf. Fig. 4C. The corresponding genotypic configuration is shown in the second row of the inset. The color coding of the mutation states of the individual loci follows that of the three regions depicted in Fig. 4B. Thus at step 10 the sites colored blue belong to *ℒ*_*q*_, while those in orange constitute 𝓂_*q*_. Since *κ* = 1, the region *ℛ*_*q*_ is empty. Therefore, the only available fitness increasing transitions at this step are 0 → 1 and must be selected from the loci colored blue.

At step 10 the locus *i* = 2 highlighted by the green box undergoes a mutation leading to the configuration at step 11. As a result, the mutation number increases by one, the pointers *𝒳* − *qv* and *𝒳* − *qv* + *v* decrease by *v*, and hence the assignment of the loci into the three regions changes (cf. inset of Fig. 4C). Now the only available fitness increasing transitions are from the *ℛ*-loci shown in red and they must remove a mutation. In the example shown, the site selected next is *j* = 16 and it is highlighted by the green box. The mutation number now decreases by one and we reach the genotype at step 12.

Observe that the net effect of going from step 10 to 12 has been the *exchange* of the mutation states of loci *i* = 2 and *j* = 16, while the mutation number remains the same. This is an example of exchange compensation. From Eq. (3) we see that, because the resistance *v*_***σ***_ = *qv* has not changed, the net change in log-fitness is equal to the change in null-fitness, Δ*F*_***σ***_(*χ*) = −Δ*u*_***σ***_. Since we are considering adaptive walks, the change in log-fitness must be positive and therefore necessarily implies an increase of null-fitness.

The adaptive walk in the fluctuating phase continues by the exchange of mutation states described above. As a result, the number of loci with mutation state 1 in 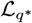 increases, terminating when all its *k* sites are mutated and leading to the fitness peak configuration shown in step 18 of Fig. 4C. Consequently, once a fitness peak has been reached, all sites *i* = 1, 2, …, *k* must be mutated, while the remaining sites carrying a mutation are selected from the subset 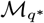, cf. Fig. 4(B), and therefore characterize the individual fitness peaks. From Eq. (9) it follows that *u*_1_ *< u*_2_ *<* … *< u*_*L*_. Consequently, the sites *i* = 1, 2, …, *k* have the lowest null-fitness. Thus the null-fitness of a fitness peak reached has two contributions: a fixed part, ∑_1≤ *i* ≤*k*_ *u*_*i*_, which is common to all fitness peaks, and a fluctuating part due to the selection of the remaining mutations from 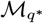. The res ulting null-fitness will therefore generally be larger than consistent with the systematic deviation of asymptotic null-fitnesses from the lower bound in Fig. 3(B).

The case of general *κ* is similar. Once the fluctuating regime is entered, the mutation number *q* performs a random walk with reflecting boundaries at *q* = *n* − *κ* + 1 and *q* = *n*, cf. Fig. 4A. Observing the genotype every time the mutation number becomes *q*\*, the change in log-fitness Δ*F*_***σ***_ from one excursion to the next is simply Δ*F*_***σ***_ = − Δ*u*_***σ***_ and hence the null-fitness continues to increase until a fitness peak is found and the walk terminates. Since the mutation number is *q*\* at the end of each excursion, the net effect on the genotype is a change of mutated loci.

### D. Effects of mutation supply and environmental fluctuation

We have so far restricted ourselves to evolutionary dynamics in the regime of low mutation rates and fixed *χ*. This has enabled us to derive analytical results and perform efficient simulations on large landscapes. However, it is essential to test the robustness of the results to effects that are present in more realistic scenarios, particularly when we relax the assumption of strong selection and weak mutations. To do this, we simulated evolution on a smaller landscape (*L* = 20) through Wright-Fisher dynamics at constant population size *N* = 10^6^ and with a mutation rate *µ* per individual and generation. We tuned the mutation rate to span four orders of magnitude and plotted the results for the null-fitness and resistance evolution in Fig. 5. At low mutation supply rates (*µN* ≈ 0.01) the results approach those of the Kimura adaptive walk, as expected. Consistent with Fig. 3A, the compensatory effect is weaker than for *L* = 100 due to smaller genotype size. As the mutation supply is increased, the compensatory phase becomes progressively less pronounced until it disappears above *µN* ≈ 1.0. Therefore, for the population size simulated here (which is similar to effective population sizes in microbial evolution experiments), biphasic adaptation occurs when *µ* is lower than ~ 10^−6^, *i*.*e*. the mutation rate per locus^2^ per generation is lower than ~ 5 × 10^−8^. For comparison, the spontaneous per base-pair mutation rates reported for *Escherichia coli* are usually in the range 10^−9^ to 10^−10^ [39–41].

**Fig 5.**
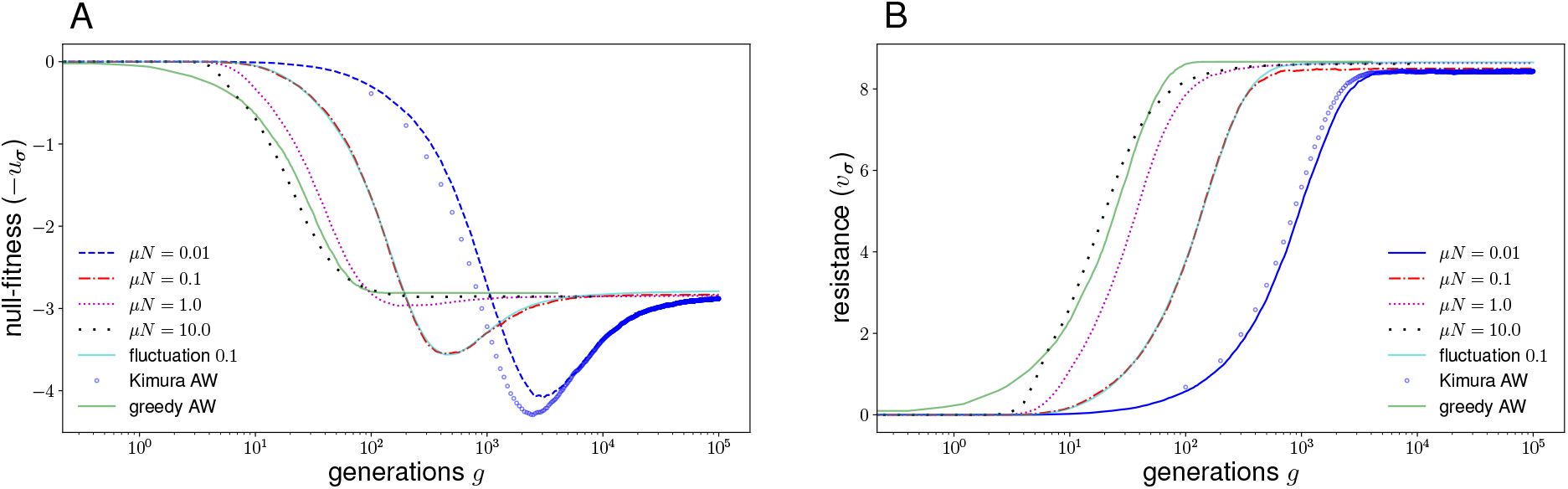
**Evolution at finite mutation rates**. The mean null-fitness (A) and resistance (B) are plotted as a function of the number of generations. The non-solid lines are the results of Wright-Fisher simulations averaged over 10^3^ landscape realizations at different mutation rates for *L* = 20 and *N* = 10^6^ at 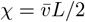; the landscape parameters are the same as in Fig 2. The solid cyan lines show mean values computed over 10^4^ landscape realizations where the stress parameter *χ* fluctuates independently in each generation according to a normal distribution with mean 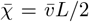 and standard deviation 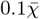 (the mutation rate is *µN* = 0.1). The blue circles show results for the Kimura adaptive walk (averaged over 10^3^ realizations), where we have multiplied the time steps by 100 to facilitate comparison withe case case *µN* = 0.01, in which one mutation occurs every 100 generations on average. The solid gray lines are the results of the greedy adaptive walk. Here the number of time steps of the walk is divided by 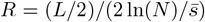, where 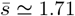 is the mean selection coefficient of mutations in the wild type background. The factor *L/*2 adjusts for the fact that on average *L/*2 mutations are required by the greedy walk algorithm to find the next mutation to be fixed (see SI), and the factor 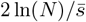 is the fixation time of a beneficial mutation with selection coefficient 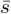 in the Wright-Fisher model. This rescaling facilitates comparison with the results from the simulations with finite mutation rates.

At high mutation rates, the population reaches slightly higher null-fitness and resistance values and on a much faster time-scale. There are two possible factors that can contribute to this effect. The first is that multiple mutants produced from an evolving genotype coexist and compete, which eventually leads to the fixation of the fittest variant in the landscape neighborhood. This is akin to the ‘greedy dynamics’ implemented in some studies on fitness landscapes [36, 42–45]. The second possible contributor is stochastic tunneling [46, 47], where the population escapes a local fitness maximum by populating neighboring lower-fitness genotypes that have evolutionary access to maxima of even higher fitness. Figure 5 shows that evolution is faster at higher mutation rates even in the directed phase, where the greedy dynamics is the only possible contributor. We conjecture that it is the dominant effect throughout the dynamics. We simulated a greedy adaptive walk where a population evolves by always moving to the fittest genotypic neighbor (see SI for further details). The curves from the greedy walk (gray lines in Fig. 5A and B) are similar to those from the Wright-Fisher model at mutation rates higher than *µN* = 1.0 in Fig. 5. In particular, a strong greedy effect eliminates the compensatory phase in both cases, since selection is competent at finding short paths to fitness maxima^3^. Our mutation supply belongs to the regime where the greedy effect was found to be dominant in an earlier theoretical study [48], consistent with our hypothesis that an efficient exploration of genotypic neighborhoods lies behind the rapid adaptation at higher mutation rates in our model.

To further test the robustness of our model, we relaxed the assumption that the environmental parameter *𝒳* is constant. In real-world scenarios, organisms frequently evolve in non-constant environments. For example, the drug concentration experienced by a microbe may vary in time due to a multitude of factors [33, 49–51]. Indeed, it is doubtful that organisms ever experience a strictly constant environment, and therefore the validity of our results depend on them being robust at least to small fluctuations in *𝒳*. To implement this, we simulated dynamics in the Wright-Fisher model where, at every generation, the value of *𝒳* was drawn independently from a normal distribution centered around a mean value of 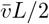. As seen from the cyan lines in Fig 5A and B, a 10% fluctuation in *χ* has negligible effect on the evolution of the phenotypes, and therefore our conclusions about the biphasic nature of evolution continue to hold.

## II. DISCUSSION

In this work, we have investigated how evolution through natural selection in a stressful environment proceeds under universal but variable antagonistic pleiotropy between null-fitness and stress resistance. We have identified two phases of adaptation, where the first phase involves resistance acquisition at the cost of null-fitness, and the second phase exhibits a partial recovery of the cost through a process we have called exchange compensation. Previous work has discussed the possibility of compensatory evolution occurring through a ‘replacement’ process, where high-cost mutants are replaced by equally resistant low-cost mutants, either by allelic replacement or mutations at different loci [14, 52]. Allelic replacement can occur through mutations that do not involve any intermediate low-resistance genotypes, but the nature of the hypothesized non-allelic replacement process has not been clarified. Exchange compensation is a specific mode of multi-step replacement that includes both reversions and fixation of new mutations. It involves individual mutational steps that can temporarily increase the cost or reduce the resistance, contrary to standard expectations from a compensatory process. Despite this, exchange compensation does not require crossing of any fitness valleys, because for every mutation that is fixed, the negative fitness effect of increased cost (or reduced resistance) is more than balanced by the increased resistance (or reduced cost) conferred by it. Rather than crossing fitness valleys [46, 47, 53], the adaptive evolution in the fluctuating regime proceeds along low dimensional ridges of the fitness landscape, the dimension of which is (in the case of the *q*-TIL model) bounded by the parameter *κ*.

A key element of our analysis is the nonlinear phenotype-fitness map (3), which is responsible for sign epistatic effects that cause the reversion of previously fixed resistance mutations in intermediate steps of the exchange compensation process. Previous work has employed phenotype-fitness maps to understand features of resistance and cost evolution. A theoretical treatment based on a two-dimensional trait space has shown how the assumption of two distinct phenotypic optima corresponding to absence and presence of antimicrobial drug can describe different scenarios of microbial adaptation to drugs [20], and some of the resulting predictions have been confirmed experimentally [22, 54]. Our study goes beyond these abstract models by using phenotypic traits with a clear biological meaning that are mapped to fitness through the empirically validated response curves (1). Furthermore, our work illustrates how epistasis in the genotype-fitness map separates the dynamics into distinct phases.

We have used particular choices for the response curve and the distribution of phenotypic effects to illustrate our results. However, the biphasic nature of evolution found here does not depend on these specific choices. The same qualitative features should be observed as long as two crucial requirements are satisfied. The first is that the dose-response curve should be such that when the resistance is sufficiently low, the dynamics should not involve reversal of resistance mutations even when they incur a high cost. This ensures the occurrence of the initial directed phase of evolution. It is shown in SI Text that this requirement is satisfied by a broad class of response curves. The second requirement is that there should be variable tradeoffs, which is crucial to exchange compensation. This requirement is not a restrictive one either, at least in the context of drug resistance, since empirical studies across microbial systems do not commonly find a strong relationship between individual mutational effects on resistance and their cost [9, 14, 55, 56]. Therefore, exchange compensation is not restricted to specific microbes or resistance mechanisms, but should occur whenever there are adaptational tradeoffs that satisfy the above-mentioned requirements.

Since exchange compensation is a slow process, its adverse effects will be most pronounced under long-term stress exposure. This has potentially important implications for drug resistance evolution. In line with the views of other authors on drug resistance[12, 57, 58], our work points to the need for better understanding of the optimal duration of a drug course. Since the cost of resistance mutations drives their reversion in the absence of the drug [9], the reduction of the cost also reduces the likelihood for drug susceptibility to be restored after treatment. Of course, real-world scenarios of drug resistance can be more complicated due to other factors not considered in this theoretical study, such as the occurrence of compensatory mutations with no adverse effects on resistance, or complex spatiotemporal variations of drug concentration in a patient [33]. While further work is needed to elucidate the role of additional factors that may complicate our modeling, the predicted generic nature of exchange compensation indicates the need for closer empirical and clinical scrutiny of the dynamics of compensatory evolution and the spread of resistance strains under long-term drug therapy.

## IV. METHODS

### A. Parameter choices

For the phenotypic effects of mutations, we use the joint distribution

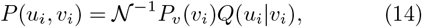

where

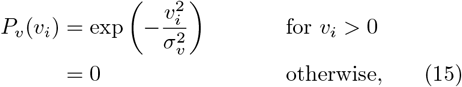

and

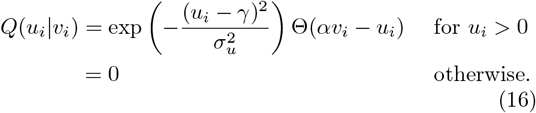

In the above, Θ is the Heaviside function and *𝒩* is a normalization constant. In short, for each mutation, we choose *v*_*i*_ from a half-normal distribution, and then choose *u*_*i*_ from a truncated normal distribution where the factor Θ(*αv*_*i*_ − *u*_*i*_) ensures that the constraint Eq. (4) is obeyed. The parameter *γ* ≥ 0 controls the mean cost of the resistance mutations. The distribution used here is slightly different from the one in [30] where the model was first introduced. The choice is made for the mathematical convenience of using variables and parameters on the logarithmic scale for a large landscape, and also in order to allow for a tunable cost of resistance.

Throughout the paper, the simulations were done with parameter values *α* = 2 and *σ*_*v*_ = *σ*_*u*_ = 1. Other parameter values are mentioned in the text and the figure captions as needed.

### B. Time-scale of adaptive walks and Wright-Fisher simulations

The algorithms for the adaptive walks and the Wright-Fisher simulations are provided in the SI and the codes are available at https://github.com/suman-g-das/Biphasic_adaptive_evo_amr. A description of Kimura and uniform adaptive walks is given in the Results section. Here we note briefly that for greedy adaptive walks (Fig. 5), each time-step consisted of randomly drawing a mutation and updating the current genotype only if the mutant was the fittest among all single mutants. Therefore, a greedy walk takes on average *L/*2 steps to find the mutation that is fixed.

For the adaptive walks, the number of time steps was identical to the number of mutations that had occurred, regardless of whether they were fixed. Moreover, origin and fixation occurred in the same step. In the Wright-Fisher simulations, the dynamics consisted of non-overlapping generations and several generations are typically needed for the fixation of a mutation. When the mutation supply rate *µN* is sufficiently small, on average 1*/µN* generations of the Wright-Fisher model are equivalent to one time-step of the Kimura adaptive walk (see Fig 5).

## ACKNOWLEDGMENTS

This work was supported by Deutsche Forschungsgemeinschaft (DFG) within CRC 1310 “Predictability in Evolution”. MM was also supported by the DFG under Projektnummer 398962893.

## Supplementary Information (SI)

### A. Mutation number in fitness maxima

As noted in the main text, there is a typical number of mutations in fitness maxima that depends on

*χ*. We provide an intuitive argument for understanding this. The log-fitness *F* is given by

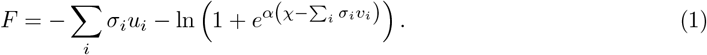

For large *χ*, the number of mutations *q* in a high fitness genotype is large, and as a crude approximation we can make the replacements 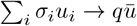 and 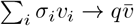, where 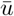 and 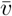 are averages over the full set of *L* mutations. This gives us the approximation

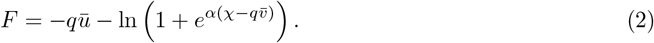

It is now straightforward to show that the leading order approximation to *q* at which the above expression is maximized is given by 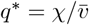.

### B. Adaptive walk simulations

Adaptive walks are discrete-time Markov chains. At any given time step, the state of the system is given by a single genotype. The simulation begins with the wild type and evolves according to the following algorithm.

Step 1. Check if the genotype is a local fitness maximum by comparing its fitness with all genotypes that differ at one locus. If yes, terminate the walk. If no, go to the next step.

Step 2. Randomly choose a genotype that differs at one locus from the current genotype.

Step 3: Update the current genotype to the chosen genotype with probability *p*. If an update occurs, go to Step 1. If not, go back to Step 2.

The probability *p* depends on the choice of adaptive walk. For the Kimura adaptive walk, *p* is given by the Kimura fixation probability (see Results section). For the uniform adaptive walk, *p* = 1 if the chosen genotype is fitter than the current one, and *p* = 0 otherwise. For the greedy adaptive walk, *p* = 1 if the chosen genotype is the fittest among all possible choices, and *p* = 0 otherwise.

### C. Sharp transition between phases

The transition from the directed to the fluctuating phase is *sharp* in the sense that it occurs on a time-scale that is much faster than the time-scales of the individual phases. To show this, we focus on the mean fixation probability of the beneficial mutations that can occur on the evolving genotype. This quantity is plotted in Figure 2A for system sizes *L* = 100, 200 as a function of the rescaled time *t/L*. The curves collapse in the directed and fluctuating phases, but not in the intermediate transition region, where the curve is steeper for larger system size. This indicates that the time-scale associated with the individual phases is *O*(*L*), but the transition occurs on faster scale. In Fig 2B, we show that rescaling the time axis by 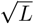 (after an appropriate shift of origin by *a*_*L*_) leads to the steep part of the curves (representing the transition) collapsing on each other. This implies that the time-scale of the transition is 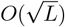, which is much shorter than the time-scale of the walk.

### D. Wright-Fisher simulations

The system evolves with non-overlapping generations at constant population size *N*. Every generation consists of a selection step and a mutation step. In the selection step, the number of offspring of the genotypic subpopulations are chosen from a multinomial distribution with probabilities proportional to their fitnesses. In the mutation step, we choose the number of mutants among the offspring of a genotypic subpopulation from a binomial distribution with parameter *µ* and then assign the offspring uniformly at random among the single-mutant genotypic neighbors of the parent genotype.

### E. Shape of response curves and existence of the directed phase

We determine the conditions under which adaptive walks starting from the wild type will display an initial phase containing only forward mutations when *𝒳* is sufficiently high. Consider a response curve (in un-transformed variables) of the form

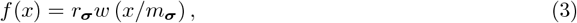

where *w*(·) is a continuous, monotonic decreasing function that goes to zero as *x* → ∞. Converting to logarithmic scale, the log-fitness is

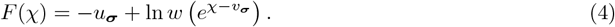

We will find *conditions on the shape of w*(·) *which ensure that, given any mutation* (*u*_*i*_, *v*_*i*_) *which is beneficial at* some *𝒳, the reversal of it must be deleterious if 𝒳 is sufficiently high*. We show below that a sufficient condition for this to happen is that the DRCs of a genotype and its single-mutation neighbor intersect at most once, *i*.*e*. the fitness difference of the two mutational neighbors cross the *χ*-axis at most once.

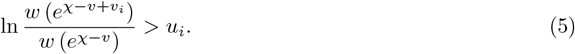

A forward mutation (*u*_*i*_, *v*_*i*_) is beneficial in a background of resistance level *v* at stress *𝒳* if and only if

The inequality is violated at *𝒳* =−∞ (corresponding to *x* = 0). Now, if the DRC’s of the background genotype and the mutant can intersect at most once, then the two sides of (5) are equal at most at one value of *𝒳* (call it 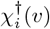). This implies that for any given values of (*u*_*i*_, *v*_*i*_) and *v*, (5) can be satisfied if and only

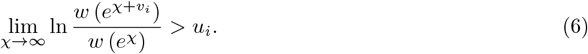

If the condition is not satisfied, the mutation is not beneficial at any *χ* in any background; if it is satisfied, then the mutation is beneficial (or equivalently, its reversal is deleterious) at sufficiently high stress (*i*.*e*., when 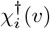). This proves the desired result, namely that a sufficient condition for the existence of a directed phase is that the response curves of mutational neighbors can intersect at most once. It was shown in [**?**] that this condition is satisfied by most functions *w*(·) used to model dose-response curves of bacteria growing in antibiotics.

Note that (6) holds for the Hill function *w*(*x*) = (1 + *x*^*α*^)^−1^ when *u*_*i*_ *< αv*_*i*_, as shown in the main text. Further, the condition holds without any restriction on the relationship between *u*_*i*_ and *v*_*i*_ when the slope of − ln *W* (*𝒳*) (where *W* (*𝒳*) = *w*(*e*^*χ*^)) increases without bound as *𝒳* → ∞. This is true when the asymptotic decay of *w*(*x*) is faster than a power law (e.g., when *w*(*x*) ~ exp(−*x*^*a*^) for any *a >* 0).

**Figure S1.**
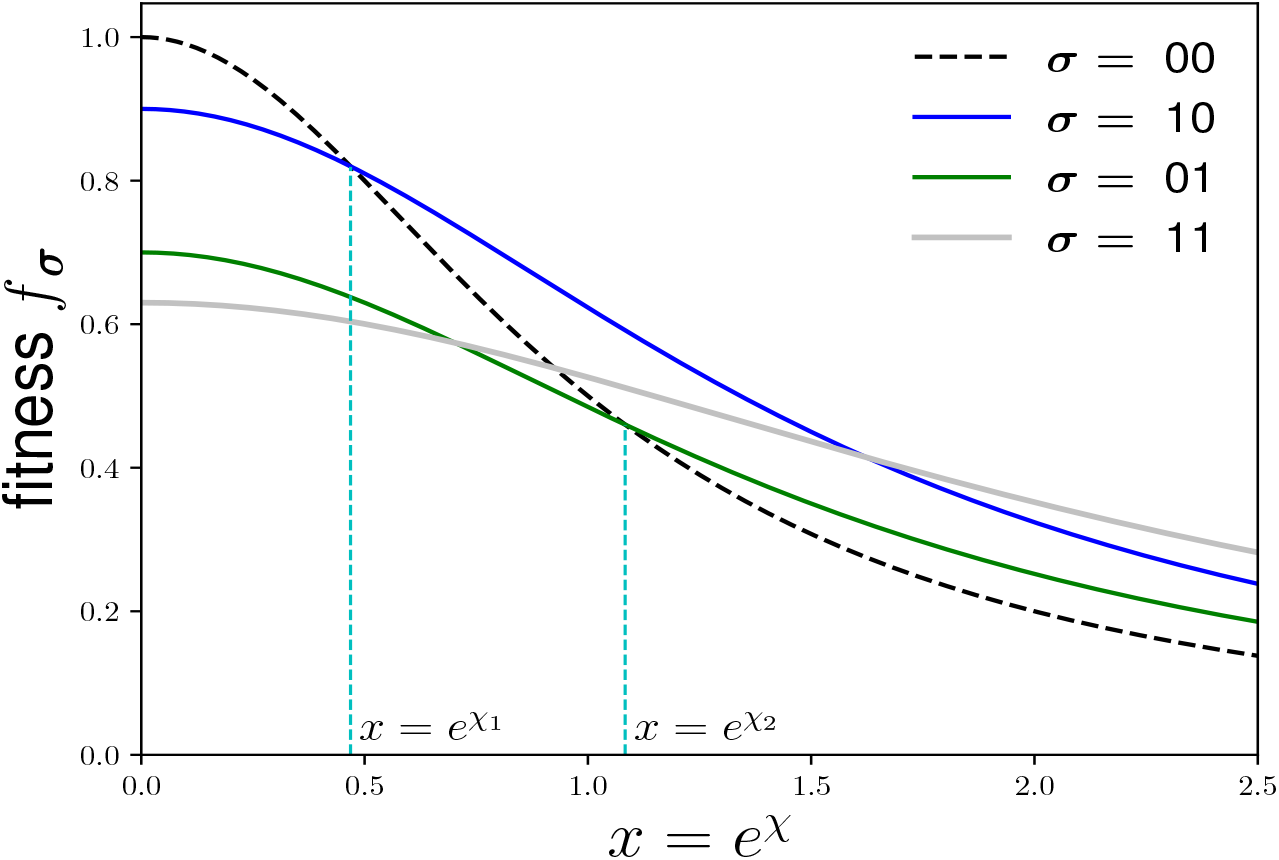
Example response curves of a two-locus system, following the Hill function form given in the main text Equation 1, with parameters *r*_1_ = 0.9, *r*_2_ = 0.7, and *m*_1_ = *m*_2_ = 1.5. The transformation of the parameters and variables between linear and log scales is given in Equation 3 of the main paper and the text immediately preceding it. The intersection points of the response curves occur at the *x*-values at which the ranks of genotypes are interchanged. For example, the single mutant 10 is fitter than the wild type 00 for *x >* exp(*𝒳* _1_).

**Figure S2.**
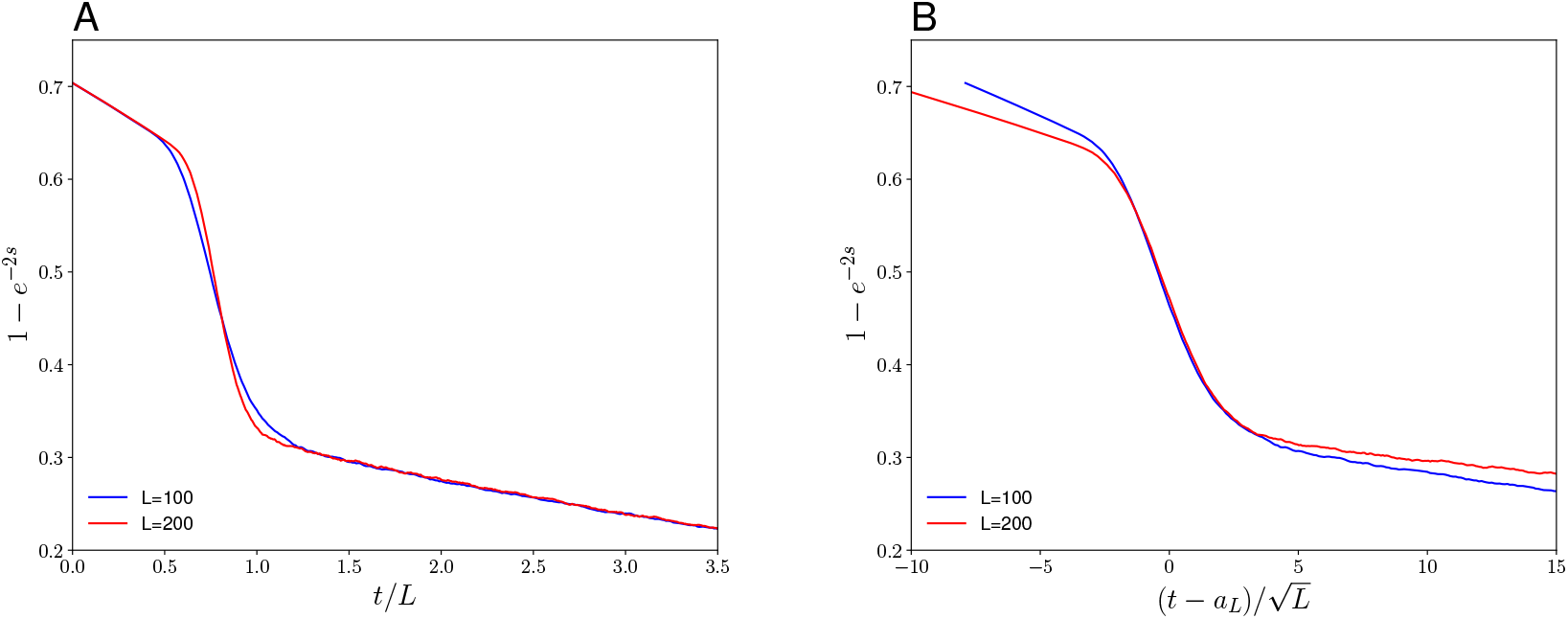
(A) The mean fixation probability of fitter neighbors of the evolving genotype as a function of time is shown for two different system sizes. The stress parameters for both sizes are 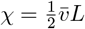. On the x-axis, *t* is the number of time steps, as defined in the main text. The time axis has been rescaled by *L* for both of the curves. (B) The same data is shown, except that the curves have been shifted along the time-axis by *a*_100_ = 79 and *a*_200_ = 158, respevc<u>ti</u>vely, which were chosen to center the curves around zero. The time axis has now been rescaled by 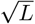.

**Figure S3.**
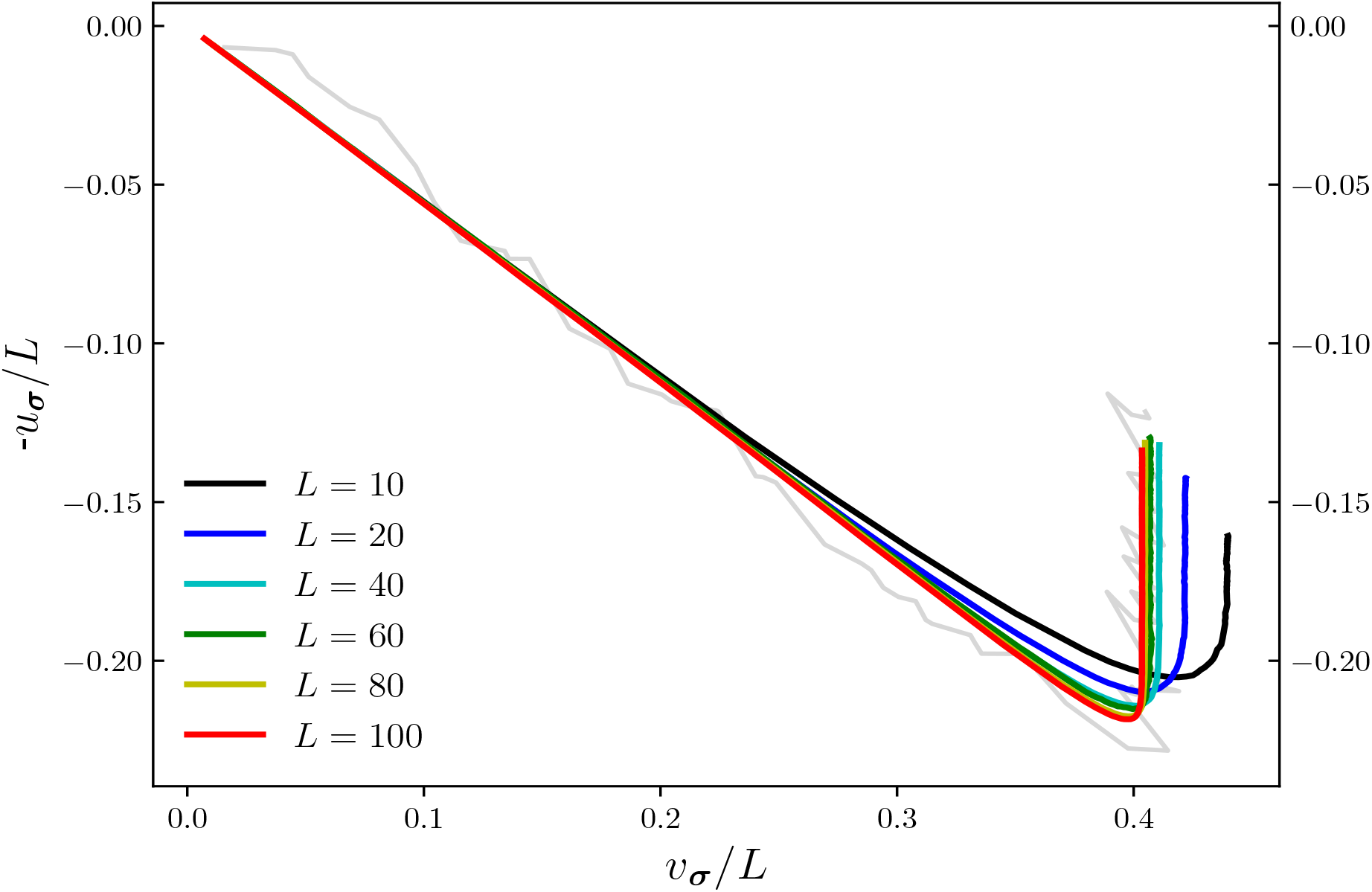
The solid curves show the mean null fitness versus the mean resistance for Kimura adaptive walks for various genome sizes. The gray line shows a single trajectory for *L* = 100. The distribution used in the plots is the given in the Methods section of the main text, with the parameter *µ* = 0. The axes have been scaled with system size *L* to facilitate comparison of the relative degree of null-fitness compensation across system sizes. The compensation increases with system size, since for larger sizes, values of *u*_*i*_ closer to zero occur with higher probability in the landscape. The degree of compensation appears to saturate at large system sizes.

## Footnotes

1 In the following we will be interested in genotypes that are neither the WT nor the all-mutant so that the sets *I*^±^ satisfying the above inequalities are non-empty.

2 In this context it is important to note that the genotype size *L* in the TIL-model represents the effective number of mutational loci that confer resistance to a specific antibiotic, rather than the total genome size.

3 For the *q*-TIL model it can be shown that the fluctuating phase disappears when greedy adaptive walks are considered.

## Notes

https://github.com/suman-g-das/Biphasic_adaptive_evo_amr

### Competing Interest Statement

The authors have declared no competing interest.

### Summary of Updates

New material, including text, tables and figures, have been added. The new version focuses more on the general lessons about compensatory evolution mediated by epistasis rather than its specific applications.

## References

[1] M. Kimura, The role of compensatory neutral mutations in molecular evolution, Journal of Genetics 64, 7 (1985).

[2] F. B.-G. Moore, D. E. Rozen, and R. E. Lenski, Pervasive compensatory adaptation in Escherichia coli, Proceedings of the Royal Society of London. Series B: Biological Sciences 267, 515 (2000).

[3] J. Bjorkman, I. Nagaev, O. Berg, D. Hughes, and D. I. Andersson, Effects of environment on compensatory mutations to ameliorate costs of antibiotic resistance, Science 287, 1479 (2000).

[4] S. Maisnier-Patin and D. I. Andersson, Adaptation to the deleterious effects of antimicrobial drug resistance mutations by compensatory evolution, Research in microbiology 155, 360 (2004).

[5] D. M. Weinreich, R. A. Watson, and L. Chao, Perspective: Sign epistasis and genetic costraint on evolutionary trajectories, Evolution 59, 1165 (2005).

[6] B. Szamecz, G. Boross, D. Kalapis, K. Kovács, G. Fekete, Z. Farkas, V. Lázár, M. Hrtyan, P. Kemmeren, M. J. Groot Koerkamp, et al., The genomic landscape of compensatory evolution, PLoS biology 12, e1001935 (2014).

[7] D. N. Ivankov, A. V. Finkelstein, and F. A. Kondrashov, A structural perspective of compensatory evolution, Current opinion in structural biology 26, 104 (2014).

[8] A. H. Melnyk, A. Wong, and R. Kassen, The fitness costs of antibiotic resistance mutations, Evolutionary applications 8, 273 (2015).

[9] D. I. Andersson and D. Hughes, Antibiotic resistance and its cost: is it possible to reverse resistance?, Nature Reviews Microbiology 8, 260 (2010).

[10] K. Schmidlin, S. Apodaca, D. Newell, A. Sastokas, G. Kinsler, and K. Geiler-Samerotte, Distinguishing mutants that resist drugs via different mechanisms by examining fitness tradeoffs, eLife 13, RP94144 (2024).

[11] A. Hinz, A. Amado, R. Kassen, C. Bank, and A. Wong, Unpredictability of the fitness effects of antimicrobial resistance mutations across environments in Escherichia coli, Mol. Biol. Evol. 41, msae086 (2024).

[12] P. Durão, R. Balbontín, and I. Gordo, Evolutionary mechanisms shaping the maintenance of antibiotic resistance, Trends in microbiology 26, 677 (2018).

[13] B. R. Levin, V. Perrot, and N. Walker, Compensatory mutations, antibiotic resistance and the population genetics of adaptive evolution in bacteria, Genetics 154, 985 (2000).

[14] M. G. Reynolds, Compensatory evolution in rifampin-resistant Escherichia coli, Genetics 156, 1471 (2000).

[15] A. K. A. Emane, X. Guo, H. E. Takiff, and S. Liu, Drug resistance, fitness and compensatory mutations in Mycobacterium tuberculosis, Tuberculosis 129, 102091 (2021).

[16] I. Nagaev, J. Björkman, D. I. Andersson, and D. Hughes, Biological cost and compensatory evolution in fusidic acid-resistant staphylococcus aureus, Molecular microbiology 40, 433 (2001).

[17] A. Handel, R. R. Regoes, and R. Antia, The role of compensatory mutations in the emergence of drug resistance, PLoS computational biology 2, e137 (2006).

[18] P. Schulz zur Wiesch, J. Engelstadter, and S. Bonhoeffer, Compensation of fitness costs and reversibility of antibiotic resistance mutations, Antimicrobial agents and chemotherapy 54, 2085 (2010).

[19] M. Baym, T. D. Lieberman, E. D. Kelsic, R. Chait, R. Gross, I. Yelin, and R. Kishony, Spatiotemporal microbial evolution on antibiotic landscapes, Science 353, 1147 (2016).

[20] T. Lenormand, N. Harmand, and R. Gallet, Cost of resistance: an unreasonably expensive concept, Rethinking Ecology 3, 51 (2018).

[21] W. Paulander, S. Maisnier-Patin, and D. I. Andersson, Multiple mechanisms to ameliorate the fitness burden of mupirocin resistance in Salmonella typhimurium, Molecular microbiology 64, 1038 (2007).

[22] N. Harmand, R. Gallet, G. Martin, and T. Lenormand, Evolution of bacteria specialization along an antibiotic dose gradient, Evolution Letters 2, 221 (2018).

[23] D. M. Weinreich, N. F. Delaney, M. A. DePristo, and D. L. Hartl, Darwinian evolution can follow only very few mutational paths to fitter proteins, Science 312, 111 (2006).

[24] M. F. Schenk, I. G. Szendro, M. L. M. Salverda, J. Krug, and J. A. G. M. de Visser, Patterns of epistasis between beneficial mutations in an antibiotic resistance gene, Molecular biology and evolution 30, 1779 (2013).

[25] P. M. Mira, J. C. Meza, A. Nandipati, and M. Barlow, Adaptive landscapes of resistance genes change as antibiotic concentrations change, Molecular biology and evolution 32, 2707 (2015).

[26] C. B. Ogbunugafor, C. S. Wylie, I. Diakite, D. M. Weinreich, and D. L. Hartl, Adaptive landscape by environment interactions dictate evolutionary dynamics in models of drug resistance, PLoS computational biology 12, e1004710 (2016).

[27] A. D. Farr, D. Pesce, S. G. Das, M. P. Zwart, and J. A. G. de Visser, The fitness of beta-lactamase mutants depends nonlinearly on resistance level at sublethal antibiotic concentrations, MBio 14, e00098 (2023).

[28] A. Papkou, L. Garcia-Pastor, J. A. Escudero, and A. Wagner, A rugged yet easily navigable fitness landscape, Science 382, eadh3860 (2023).

[29] J. Zhang, Patterns and evolutionary consequences of pleiotropy, Annual Review of Ecology, Evolution, and Systematics 54, 1 (2023).

[30] S. G. Das, S. O. Direito, B. Waclaw, R. J. Allen, and J. Krug, Predictable properties of fitness landscapes induced by adaptational tradeoffs, eLife 9, e55155 (2020).

[31] S. G. Das, J. Krug, and M. Mungan, Driven disordered systems approach to biological evolution in changing environments, Physical Review X 12, 031040 (2022).

[32] R. R. Regoes, C. Wiuff, R. M. Zappala, K. N. Garner, F. Baquero, and B. R. Levin, Pharmacodynamic functions: a multiparameter approach to the design of antibiotic treatment regimens, Antimicrobial agents and chemotherapy 48, 3670 (2004).

[33] E. S. King, D. S. Tadele, B. Pierce, M. Hinczewski, and J. G. Scott, Diverse mutant selection windows shape spatial heterogeneity in evolving populations, PLOS Computational Biology 20, e1011878 (2024).

[34] M. Knopp and D. I. Andersson, Predictable phenotypes of antibiotic resistance mutations, mBio 9, e00770 (2018).

[35] J. H. Gillespie, Molecular evolution over the mutational landscape, Evolution 38, 1116 (1984).

[36] S. Kauffman and S. Levin, Towards a general theory of adaptive walks on rugged landscapes, Journal of theoretical Biology 128, 11 (1987).

[37] D. M. McCandlish and A. Stoltzfus, Modeling evolution using the probability of fixation: history and implications, The Quarterly review of biology 89, 225 (2014).

[38] J. Domingo, P. Baeza-Centurion, and B. Lehner, The causes and consequences of genetic interactions (epistasis), Annu. Rev. Genom. Hum. Genet. 20, 17.1–17.28 (2019).

[39] J. W. Drake, A constant rate of spontaneous mutation in dna-based microbes., Proceedings of the National Academy of Sciences 88, 7160 (1991).

[40] S. Wielgoss, J. E. Barrick, O. Tenaillon, S. Cruveiller, B. Chane-Woon-Ming, C. Médigue, R. E. Lenski, and D. Schneider, Mutation rate inferred from synonymous substitutions in a long-term evolution experiment with escherichia coli, G3: Genes— Genomes— Genetics 1, 183 (2011).

[41] A. B. Williams, Spontaneous mutation rates come into focus in escherichia coli, DNA repair 24, 73 (2014).

[42] H. A. Orr, A minimum on the mean number of steps taken in adaptive walks, Journal of Theoretical Biology 220, 241 (2003).

[43] J. Franke and J. Krug, Evolutionary accessibility in tunably rugged fitness landscapes, Journal of Statistical Physics 148, 706 (2012).

[44] J. A. G. M. de Visser and J. Krug, Empirical fitness landscapes and the predictability of evolution, Nature Reviews Genetics 15, 480 (2014).

[45] S.-C. Park, J. Neidhart, and J. Krug, Greedy adaptive walks on a correlated fitness landscape, Journal of theoretical biology 397, 89 (2016).

[46] D. M. Weinreich and L. Chao, Rapid evolutionary escape by large populations from local fitness peaks is likely in nature, Evolution 59, 1175 (2005).

[47] Y. Iwasa, F. Michor, and M. A. Nowak, Stochastic tunnels in evolutionary dynamics, Genetics 166, 1571 (2004).

[48] I. G. Szendro, J. Franke, J. A. G. de Visser, and J. Krug, Predictability of evolution depends nonmonotonically on population size, Proceedings of the National Academy of Sciences 110, 571 (2013).

[49] D. W. Kolpin, M. Skopec, M. T. Meyer, E. T. Furlong, and S. D. Zaugg, Urban contribution of pharmaceuticals and other organic wastewater contaminants to streams during differing flow conditions, Science of the Total Environment 328, 119 (2004).

[50] E. S. King, J. Pelesko, J. Maltas, S. J. Owen, E. Dolson, and J. G. Scott, Fitness seascapes facilitate the prediction of therapy resistance under time-varying selection, bioRxiv, 2022 (2022).

[51] R. Gross, M. Mungan, S. G. Das, M. Yüksel, B. Maier, T. Bollenbach, J. Krug, and J. A. G. de Visser, Collective β-lactam resistance in Escherichia coli due to β-lactamase release upon cell death, bioRxiv, 2024 (2024).

[52] F. M. Cohan, E. C. King, and P. Zawadzki, Amelioration of the deleterious pleiotropic effects of an adaptive mutation in Bacillus subtilis, Evolution 48, 81 (1994).

[53] C. S. Gokhale, Y. Iwasa, M. A. Nowak, and A. Traulsen, The pace of evolution across fitness valleys, Journal of theoretical biology 259, 613 (2009).

[54] N. Harmand, R. Gallet, R. Jabbour-Zahab, G. Martin, and T. Lenormand, Fisher’s geometrical model and the mutational patterns of antibiotic resistance across dose gradients, Evolution 71, 23 (2017).

[55] K. M. Brown, M. S. Costanzo, W. Xu, S. Roy, E. R. Lozovsky, and D. L. Hartl, Compensatory mutations restore fitness during the evolution of dihydrofolate reductase, Mol. Biol. Evol. 27, 2682 (2010).

[56] G. Brandis, F. Pietsch, R. Alemayehu, and D. Hughes, Comprehensive phenotypic characterization of rifampicin resistance mutations in Salmonella provides insight into the evolution of resistance in Mycobacterium tuberculosis, Journal of Antimicrobial Chemotherapy 70, 680 (2015).

[57] H. Lambert, Don’t keep taking the tablets?, The Lancet 354, 943 (1999).

[58] M. J. Llewelyn, J. M. Fitzpatrick, E. Darwin, C. Gorton, J. Paul, T. E. Peto, L. Yardley, S. Hopkins, A. S. Walker, et al., The antibiotic course has had its day, Bmj 358 (2017).

